# Mechanosensory neuron regeneration in adult *Drosophila*

**DOI:** 10.1101/812057

**Authors:** Ismael Fernández-Hernández, Evan B. Marsh, Michael A. Bonaguidi

## Abstract

Auditory and vestibular mechanosensory hair cells do not regenerate following injury or aging in the adult mammalian inner ear, inducing irreversible hearing loss and balance disorders for millions of people. Research on model systems showing replacement of mechanosensory cells can provide mechanistic insights into developing new regenerative therapies. Here, we developed lineage tracing systems to reveal, for the first time, the generation of mechanosensory neurons in the Johnston’s Organ (JO) of intact adult *Drosophila*, which are the functional counterparts to hair cells in vertebrates. New JO neurons develop cilia, express an essential mechano-transducer gene and target central brain circuitry. Furthermore, we identified self-replication of JO neurons as an unexpected mechanism of neuronal plasticity, which is enhanced upon treatment with experimental and ototoxic compounds. Our findings introduce a new platform to expedite research about mechanisms and compounds mediating mechanosensory cell regeneration, with implications for hearing and balance restoration in humans.

**SUMMARY STATEMENT:** Using refined lineage tracing and live imaging, we identified self-renewal of mechanosensory neurons in adult *Drosophila*, the functional counterparts to vertebrate hair cells, and their enhanced regeneration through pharmacological administration.

## INTRODUCTION

Hearing and balance disorders affect over 5% of the world’s population, with 1 in 3 people affected by the age of 80 (Geleoc and Holt, 2014). By the year 2050, 900 million people are expected to have hearing and balance disorders (WHO 2020). These disorders are due to the degeneration of mechanosensory hair cells and their innervating neurons in the inner ear, following damage by genetic mutations, excessive noise, ototoxic drugs or aging. Unfortunately, no treatments exist to replenish lost cells in the human sensory epithelia (Müller and Barr-Gillespie, 2015). Thus, regenerative strategies are urgently needed to recover auditory and vestibular function for millions of people. While non-mammalian vertebrates are able to functionally replenish hair cells throughout life (Kniss et al., 2016; Ryals et al., 2013; Stone and Cotanche, 2007), mammals show scarce regenerative capacity in the cochlea at early postnatal stages (Bramhall et al., 2014; Cox et al., 2014; Kelley et al., 1995; White et al., 2006) and low levels in vestibular organs during adulthood (Bucks et al., 2017; Forge et al., 1993; Golub et al., 2012; Kawamoto et al., 2009; Warchol et al., 1993). In all cases, non-sensory supporting cells trans-differentiate to regenerate hair cells (Atkinson et al., 2015; Bucks et al., 2017; Kniss et al., 2016; Stone and Cotanche, 2007; White et al., 2006). Still, research on these models at the genetic, cellular, circuitry and behavioral levels is costly and technically challenging.

The fruit fly *Drosophila melanogaster* harbors ciliated mechanosensory neurons in the Johnston’s Organ (JO) on the second segment of its two antennae. JO neurons are clustered in ~200 scolopidia per antenna, comprising multicellular units with 2-3 JO neurons and surrounding supporting cells (Albert and Go, 2015; Boekhoff-Falk and Eberl, 2014; Ishikawa and Kamikouchi, 2016). JO neurons develop in response to conserved genetic programs and act as counterparts to mammalian hair cells and their innervating neurons by supporting auditory and vestibular functions (Boekhoff-Falk, 2005; Eberl and Boekhoff-Falk, 2007; Kamikouchi et al., 2009; Li et al., 2018; Sun et al., 2009; Wang et al., 2002). *Drosophila* represents a compelling platform to accelerate research on the functional regeneration of mechanosensory cells, due to: genome-wide available genetic tools; the detailed characterization of JO neurons at the circuitry and behavioral levels (Ishikawa et al., 2017; Kamikouchi et al., 2006; Lai et al., 2012; Matsuo et al., 2016; Vaughan et al., 2014); and simple scalability at low cost. Even so, turnover of JO neurons has not yet been reported. Previous reports demonstrate a proliferative capacity in the brain of adult *Drosophila*, both in physiologic conditions and following injury (Fernández-Hernández et al., 2013; Kato et al., 2009; Li et al., 2020). Therefore, we hypothesized that the peripheral nervous system also has this capacity for proliferation. To test this, we implemented multiple modified lineage tracing systems to reveal adult-born JO neurons by live imaging of intact adult *Drosophila*. We observed that new JO neurons acquire features of mature, functional neurons. We also captured the self-renewal of JO neurons at low-frequency — a newly discovered mechanism that counteracts neuronal cell death. Furthermore, the oral administration of drugs accelerates JO self-renewal within intact flies. Our results underscore the broad potential of this new *in vivo* platform for understanding and promoting mechanosensory cell regeneration.

## RESULTS

### P-MARCM detects adult neurogenesis in Drosophila brain

We previously developed a Perma-Twin system to detect low levels of cell proliferation and regeneration in the adult *Drosophila* brain (Fernández-Hernández et al., 2013). However, this method required antibodies to assess the identity of new cells (e.g., elav for neurons); also, quantification of new cells relied upon resolving cells labeled only by cytoplasmic fluorescent reporters. Further, this method did not permit the genetic manipulation of adult-born cells. In order to overcome these limitations and detect low levels of cell proliferation in adult *Drosophila* in a cell type-specific and sustained manner, we developed P-MARCM (Permanent-MARCM). Built into MARCM (Lee and Luo, 1999), P-MARCM can detect sporadic mitotic events in virtually any adult tissue, and label newly generated cell populations. To activate P-MARCM, a heat shock-induced pulse of FLIPPASE (FLP) excises an FRT-flanked STOP codon between *tubuline* promoter and *lexA* transactivator (Singh et al., 2013), which in turn binds to *lexAOp-Flp* sequence (Pfeiffer et al., 2010) to drive FLIPPASE permanently in heat shock-responding cells (Fig. 1A,B). Cell-type specificity is achieved by incorporating desired *GAL4* lines to express nuclear-localized RFP (Barolo et al., 2004) and cytoplasmic GFP (Shearin et al., 2014) only in adult-born cells of interest in a given lineage, making antibodies dispensable to assess cell identity. These enhanced fluorescent reporters allow for the concurrent assessment of detailed morphology and straight-forward cell quantification. Furthermore, the introduction of an additional UAS-transgene permits genetic manipulation of adult-born cells and assessment of their functional contributions (Fig. 1C). To benchmark the utility of P-MARCM, we used a pan-neuronal *nsyb-GAL4* line and detected previously reported physiologic neurogenesis in the adult optic lobes, where we also detected PH3+ cells, (Fig. S1), and neuronal regeneration upon injury (Fig. S2) (Fernández-Hernández et al., 2013). Therefore, P-MARCM can capture adult-cell proliferation in a cell type-specific manner under both physiological and injury conditions.

**Figure 1.**
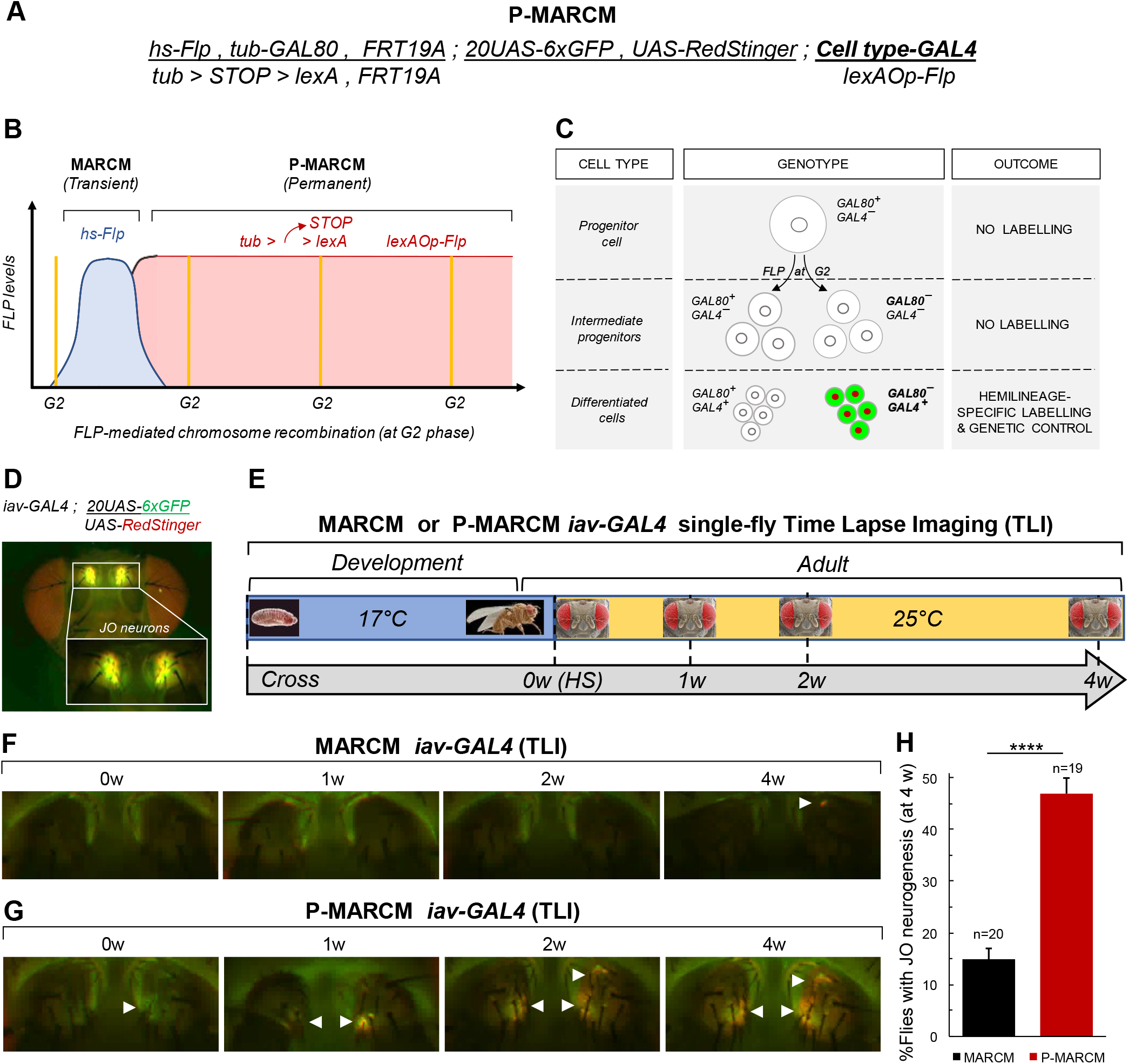
P-MARCM live imaging reveals Johnston’s Organ (JO) neurogenesis in adult *Drosophila*. (A) P-MARCM system to label and genetically manipulate adult-born cells in a cell type-specific manner. (B) P-MARCM becomes permanently active via constitutive Flippase (FLP)-mediated mitotic chromosome recombination to capture slowly dividing cells. (C) P-MARCM labels adult-born cells of interest in a lineage by cell type-specific *GAL4* lines (*iav-GAL4* for Johnston’s Organ neurons). Engineered nuclear-red and membrane-green fluorescent reporters allows quantification and cellular morphology assessment without antibodies. (D) Johnston Organ (JO) neuron visualization in live flies by Time-Lapse Imaging (TLI) using *iav-GAL4* with engineered fluorescent reporters. (E) Experimental strategy to capture JO adult neurogenesis by MARCM or P-MARCM. Flies are kept at 17°C during development to minimize leaky activation. 2-5 days-old flies are activated via heat shock and individual antennae are imaged over 4 weeks (w) by fluorescent microscopy. (F-H) Time-lapse imaging of P-MARCM identifies JO neurogenesis in *Drosophila* antennae (n=20, Fig. 1G) more frequently than MARCM (n=19, Fig. 1F) (*p=0,00014*, cumulative probability on binomial distribution: Fig 1H). Error bars represent s.e.m.

### Identification of JO neurogenesis by P-MARCM

In order to assess the generation of JO neurons in adult *Drosophila*, we used P-MARCM containing *iav-GAL4* line (Ishikawa et al., 2017; Kwon et al., 2010). *Iav* (inactive) is a transient receptor potential (TRP) vanilloid channel expressed exclusively in the cilia of chordotonal neurons, and is essential for mechano-transduction in hearing (Boekhoff-Falk and Eberl, 2014; Gong et al., 2004) (Fig. S3). Importantly, incorporation of enhanced nuclear-RFP (Barolo et al., 2004) and cytoplasmic GFP (Shearin et al., 2014) reporters in P-MARCM allowed for direct identification of JO neurons by live imaging of intact flies (Fig. 1D). We therefore activated P-MARCM *iav-GAL4* in adult flies and conducted single-fly time-lapse imaging over 4 weeks (Fig. 1E). Remarkably, this approach revealed JO neuron generation in P-MARCM flies (47%, n=19 flies) at a higher frequency than transient, regular MARCM (15%, n=20 flies) (Fig. 1F-H).

We then quantified JO neurogenesis by confocal imaging of antennae dissected from P-MARCM *iav-GAL4* flies over 4 weeks (Fig. 2A). While MARCM barely detected neurogenesis, P-MARCM captured increasing amounts of JO neurogenesis over time (Fig. 2B-D). To better account for background levels found in the control group (i.e., non-HS flies at 0w, 4.9 +/-3.9s.d. cells/fly, n=16 flies, Fig. S4A), we ran a Gaussian mixture model (Fig. S4B) and determined that any fly in experimental (i.e. HS) groups with 10 or more labelled JO neurons can be regarded as undergoing adult neurogenesis with 94% certainty. P-MARCM captured 10-36 JO neurons/fly across time points (avg. 20.2 +/-7.3s.d. neurons/fly) (Fig. 2 B-D, Fig. S4A). Furthermore, after clustering flies into “Responders” or “Non-responders” (*i.e*., JO neurogenesis present or absent, Fig. S4B) we observed a significant increase in the number of “Responders” over time (Fig. 2E), with 57% by 4 weeks and 42% across time points. This outcome is consistent with the 47% ratio detected by our live imaging approach (Fig. 1H). Taken together, our results identify JO neurogenesis in adult *Drosophila* using complementary *in vivo* time-lapse imaging and confocal microscopy approaches.

**Figure 2.**
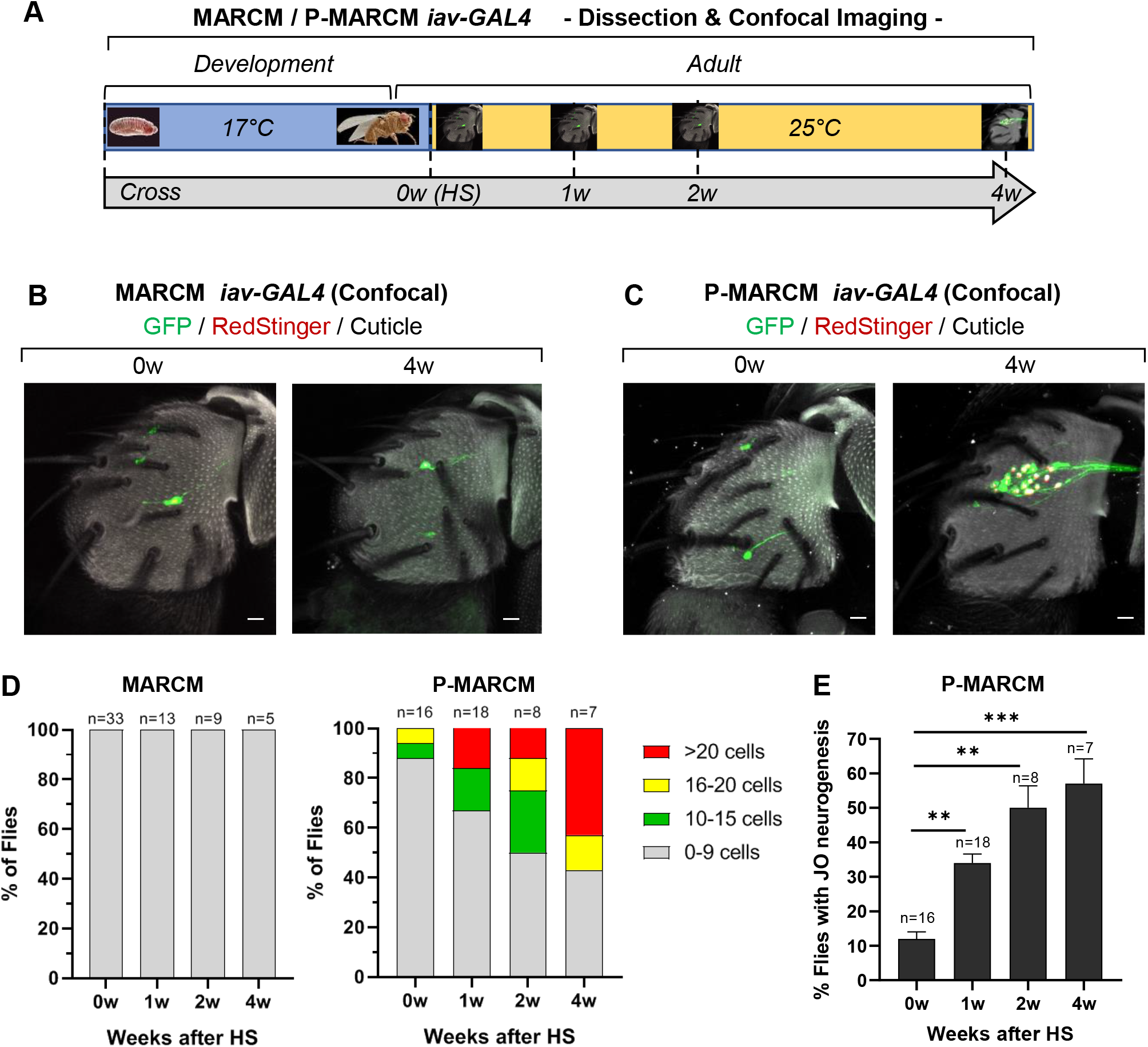
Quantification of adult JO neurogenesis. (A) Experimental strategy to compute adult JO neurogenesis using MARCM and P-MARCM *iav-GAL4* strategies. Two-5 day–old flies are Heat-Shocked (HS) to activate the MARCM or P-MARCM system and antennae are dissected up to 4 weeks (w) later for quantification. (B) Transient MARCM *iav-GAL4* does not capture JO neurogenesis. Scale bar: 10μm. (C) P-MARCM *iav-GAL4* reveals adult-born JO neurons. Scale bar: 10μm. (D) The number of adult-born JO neurons increase over time in antennae of P-MARCM, but not MARCM flies. (E) The number of flies with JO neurogenesis increases over time. \*\**p<0,01*; \*\*\**p<0,001*, cumulative probability on binomial distribution. Bars represent mean +/-s.e.m.

### Adult-born JO neurons mature and target brain circuitry

We next evaluated the cellular features of newborn JO neurons. We performed confocal analysis on antennae and brains of P-MARCM *iav-GAL4* flies with JO neurogenesis detected by live imaging by 4 weeks (Fig. 3A) (representative images shown for fly in Fig. 1G). At the cellular level, adult-born JO neurons develop cilia, express the essential mechano-transducer channel *iav* (Gong et al., 2004), and extend axons. Neuronal morphology, including cilia, was visualized in adult-born JO neurons using a cytoplasmic hexameric GFP with enhanced signal (Shearin et al., 2014) without a need for antibodies (Fig. 3B, arrowheads). At the circuit level, new JO neurons project axons to the brain through the Antennal Mechanosensory and Motor Center (AMMC) in both, auditory (high frequency, Zone A; low frequency, Zone B) and vestibular (backward deflections, Zone E) circuits (Ishikawa and Kamikouchi, 2016; Kamikouchi et al., 2006) (Fig. 3C). These features were consistently found in all cases containing JO neurogenesis. Therefore, these observations strongly suggest new JO neurons can mature and functionally remodel mechanosensory circuitry.

**Figure 3.**
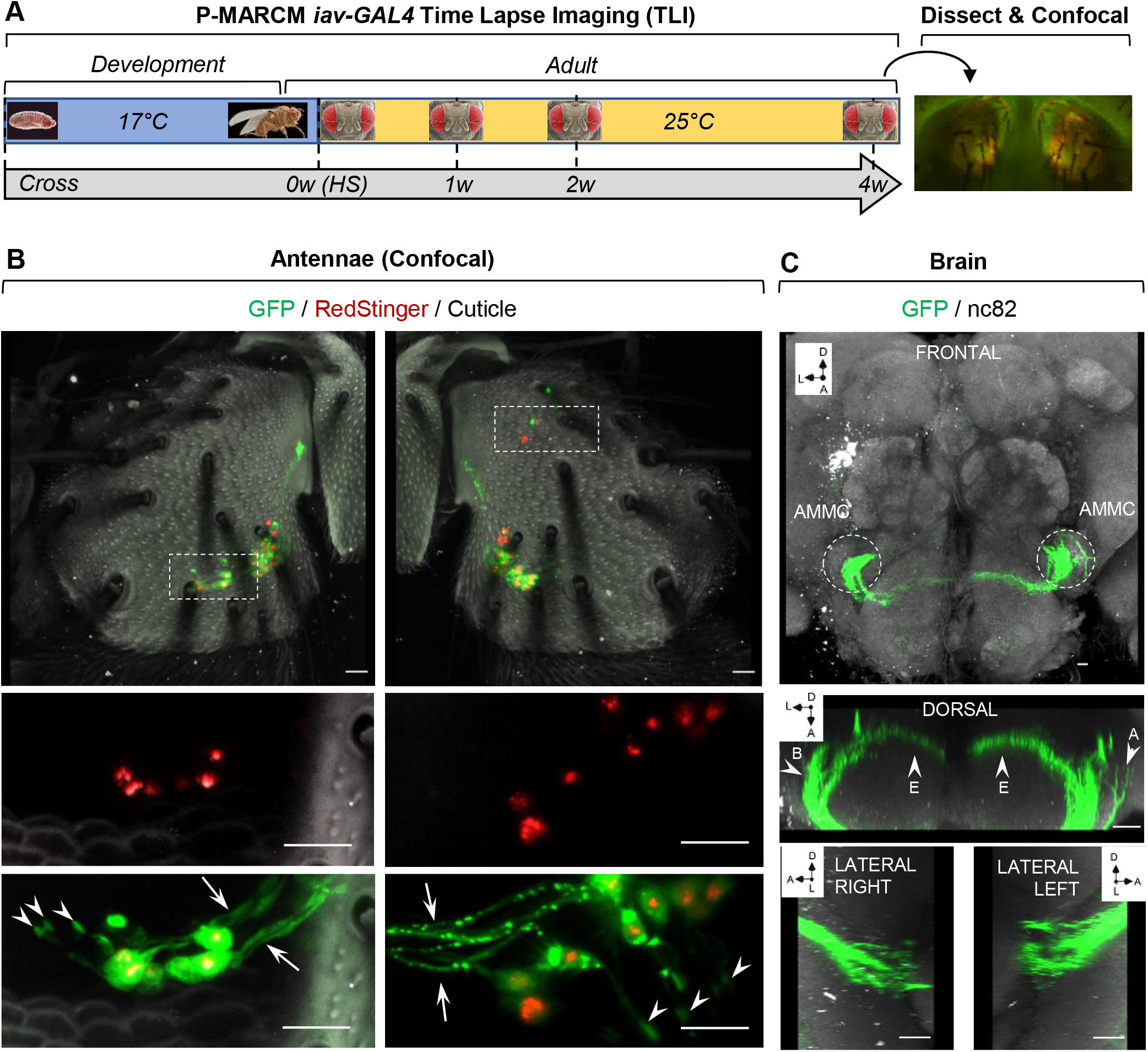
Adult-born JO neurons acquire mechanosensory features and target brain circuitry. (A) P-MARCM *iav-GAL4* flies with adult JO neurogenesis detected by Time Lapse Imaging (see also Figure 1F) were dissected for detailed cellular analysis by confocal microscopy. w: week. (B) Adult-born JO neurons develop cilia (arrowhead), express the essential mechanotransducer channel *iav* and project axons to the brain (arrows). Scale bar: 10 μm. (C) Axons from adult-born JO neurons target the brain through the Antennal Mechanosensory and Motor Center (AMMC) in both auditory (High Freq, A and Low Freq, B) and vestibular (backward deflections, E) circuit patterns. A: Anterior. D: Dorsal. L: Lateral according to Ishikawa and Kamikouchi, 2016. Scale bars for all panels: 10 μm.

### Self-replication of JO neurons

We next sought to identify a cellular source for adult-born JO neurons. In vertebrates, hair cell regeneration involves trans-differentiation of non-sensory supporting cells (Atkinson et al., 2015; Brignull et al., 2009). Recent studies demonstrate various modes of cell replacement, including proliferation of undifferentiated progenitors, de-differentiation and division of mature cells, and direct mitosis of post-mitotic cell types (Post and Clevers, 2019). We therefore considered 3 possible sources of adult JO neurons: 1) undifferentiated progenitors maintained from development; 2) non-sensory supporting cells in the scolopidium; and 3) pre-existing JO neurons. To gain insight into possible mechanisms, we observed labeled cells for features of cell division. A detailed analysis of P-MARCM-*iav* and MARCM-iav confocal images revealed instances of split DNA on single JO neurons, and pairs of labeled JO with intermingled cilia and axons (Fig. 4A-C), suggesting the possibility of JO self-division (1.3%, n=557 neurons analyzed).

**Figure 4.**
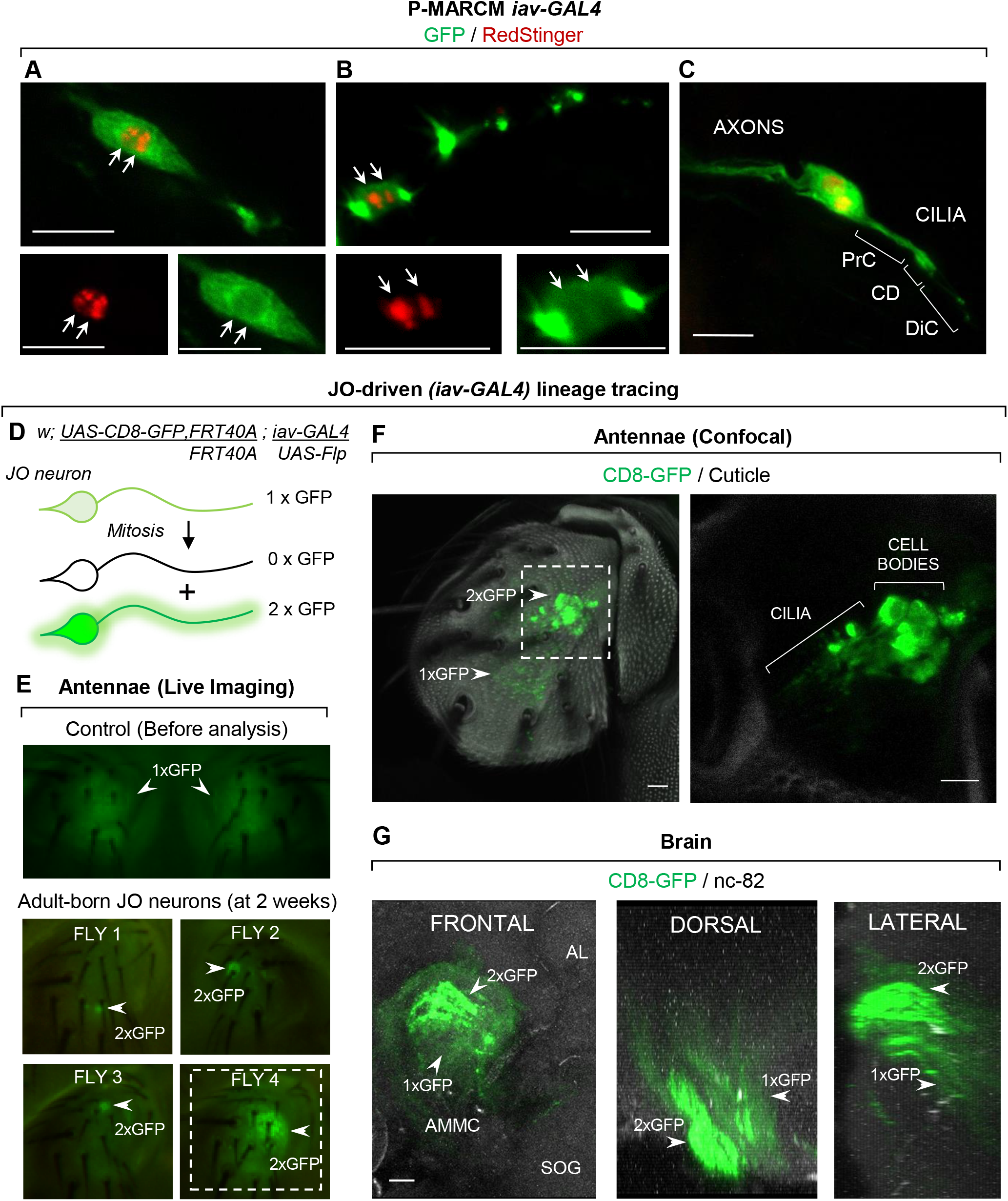
Adult JO neurons undergo self-division. (A-C) P-MARCM *iav-GAL4* captures JO neurons with split DNA (A, B) and close proximity (C), suggestive of mitotic activity. PrC: Proximal Cilia; CD: Ciliary Dilation; DiC: Distal Cilia. Scale bars: 10μm (D) A lineage tracing system to assess JO self-division: *iav-GAL4* drives constitutive expression of membrane-tethered GFP and FLP recombinase in JO neurons. Upon mitosis, one JO neuron becomes homozygous for GFP and reveals self-division. (E) JO self-division is detected by live fluorescent microscopy in flies at 2 weeks after ecclosion (arrowheads). Fly 4 is shown in panel (F). (F) A cluster of self-replicated JO neurons are detected in the antennae by confocal microscopy (G) Axons expressing two copies of GFP appear in the brain in an Auditory (Zone B) circuit pattern, indicating mature JO neuronal identity via self-division. AL: Antennal Lobe; AMMC: Antennal Mechanosensory and Motor Center; SOG: Subesophageal Ganglion. Scale bars for all panels: 10 μm.

We next conducted anti-PH3 immunostaining in an attempt to capture mitotic JO neurons. Sparse mitotic figures were found in the optic lobes, a cell-dense region of the brain where P-MARCM also captures proliferation (~40,000 total interneurons/optic lobe (Morante et al., 2011); 9.2 +/-1.0 s.e.m. PH3+ cells/optic lobe, n=26 optic lobes; Fig. S1D). However, we did not capture any PH3+ neurons in the JO (n=30 antennae; ~500 total JO neurons/antenna (Kamikouchi et al., 2006), data not shown). These results are consistent with low detection of mitotic figures even in continuously proliferating tissues, such as the adult posterior midgut (~2 PH3+ cells/gut) (Obata et al., 2018; Ren et al., 2010; Tian and Jiang, 2014).

To overcome detection limitations by immunostaining and to directly test mitotic activity in JO neurons, we implemented a JO-driven lineage tracing method. Specifically, JO neurons constitutively express the recombinase Flippase and a single copy of the UAS-CD8-GFP reporter, distal to an FRT site on the second chromosome. Upon mitosis, 1 daughter JO neuron becomes homozygous for GFP (2xGFP), clearly distinguishable from the single-copy GFP background (1xGFP) by live imaging (Fig. 4D-E). Remarkably, this approach captured low-level JO self-renewal at different points over 4 weeks in 20% of the flies analyzed (n=85 flies, 1-11 neurons/antenna) (Fig. 4E-F), in some cases at single-neuron resolution (Fig. S5B, C). These cells existed as twin spots (i.e. 2xGFP+ neuron paired with a 0xGFP neuron in a 1xGFP background, Fig. S5D), further supporting JO neuron mitotic division. New JO neurons contained cilia when visualized by a membrane-tethered GFP (Fig. 4F) and 2xGFP JO-axon bundles in the brain, indicating JO replication from cell bodies to terminal connections (Fig. 4G). Importantly, JO self-renewal was detected in both males and females (59% males, 41% females, n=17 flies). We ensured that neurons labeled appeared in adult stages by pre-screening every single fly before analysis and removing any “escapers,” labeled by leaky expression of FLIPPASE during development, which appeared at very low rates (2.3%, n=306 flies assessed). In this way, only flies lacking prior recombination were selected for analysis of adult neurogenesis (Fig. 4E).

In order to further determine mitotic activity in JO neurons, we assembled a third lineage tracing approach, which we called Perma Twin iav (PT-iav). In this mitotic recombination-based system, built into Twin Spot MARCM (Yu et al., 2009) and Perma Twin (Fernández-Hernández et al., 2013), differential expression of membrane-tethered GFP and RFP in each hemiclone occurs only in JO neurons arising from JO neuron self-division events, in an otherwise non-labelled background (Fig. 5A). Remarkably, PT-iav also identified mitosis in JO neurons, apparent when using both *in vivo* time-lapse tracing and confocal microscopy in flies 1 to 4 weeks-old (13% of flies, n=60 flies, 1-17 cells/antennae, Fig. 5B,E). Reproducing earlier findings, we detected that JO neurons develop cilia and project axons to the brain, demonstrating complete JO neuron self-replication (Fig. 5C, D and 5F, G). Cases of single JO neurons labelled in a single color may indicate only one daughter neuron survived and kept proliferating upon mitosis. Taken together, our results uncover the unexpected potential of *Drosophila* neurons to self-divide—a previously unknown mechanism for nervous system regeneration under physiological conditions.

**Figure 5.**
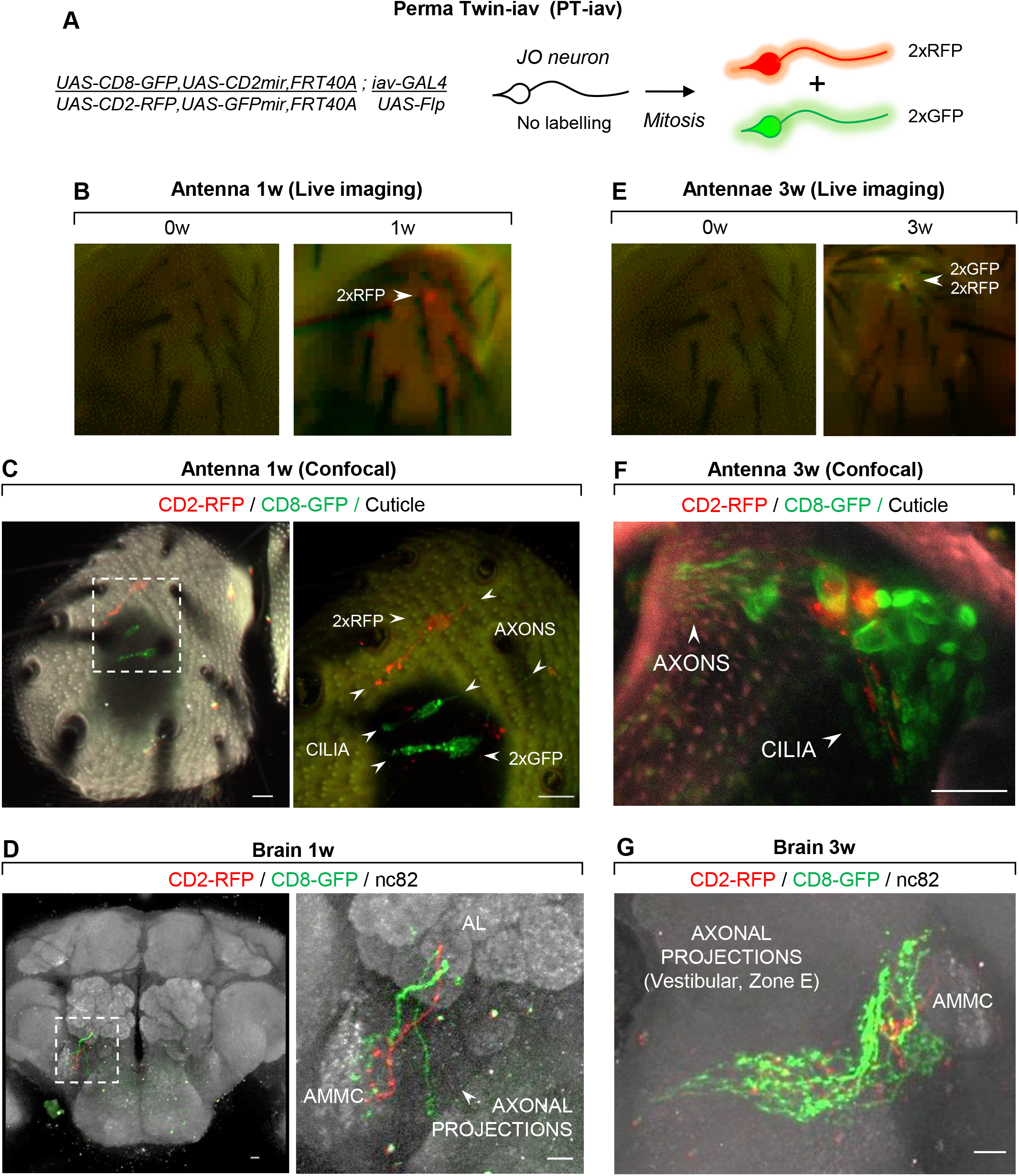
PT-iav captures JO neuron self-renewal under physiological conditions. (A) In Perma Twin-iav (PT-iav) lineage tracing, GFP and RFP reporters and repressors are differentially segregated upon JO neuron mitosis to label twin progeny hemi-clusters with GFP or RFP and reveal JO neuron self-renewal. (B) PT-iav reveals JO neuron self-renewal in 1w-old flies by live imaging. (C-D) Newborn JO neurons develop cilia and target the brain through the AMMC. (E) Self-renewed JO neurons are detected in 3w-old flies by live imaging. (F-G) New neurons develop cilia and send axonal projections to the brain in the vestibular circuit pattern (Zone E). AMMC: Antennal Mechanosensory and Motor Center; w: week. Scale bars for all panels: 10μm.

### JO cell turnover occurs under physiological conditions

Next, we asked whether JO regeneration occurs in an additive manner or as a cell turnover mechanism. Immunostaining for cleaved-Caspase3 (Ca3) revealed JO apoptosis at a higher frequency and to a greater extent in posterior neurons (5.8+/-1.0 s.e.m. Ca3+ neurons/antenna, 100% antennae, n=33), compared to anterior neurons (1.5+/-0.25 s.e.m. Ca3+ cells/antenna, 12% antennae, n=33) (Fig. 6A, C-D). Accordingly, P-MARCM captured JO neurogenesis at higher frequency and to a greater extent on posterior regions (8.7+/-2.2 s.e.m. neurons/antenna, 69% antennae, n=16), compared to anterior regions of the antenna (2.4+/-1.0 s.e.m. neurons/antenna, 44% antennae, n=16) (Fig. 6B-D). These correlations suggest JO cell turnover. Supporting this finding, the GFP-only JO-specific lineage tracing system also detected JO self-renewal in the vicinity of apoptotic cells (Fig. 6E). Taken together, these results describe cellular plasticity in the mechanosensory system of adult *Drosophila*, pointing to a cell turnover mechanism that could preserve auditory and vestibular functions.

**Figure 6.**
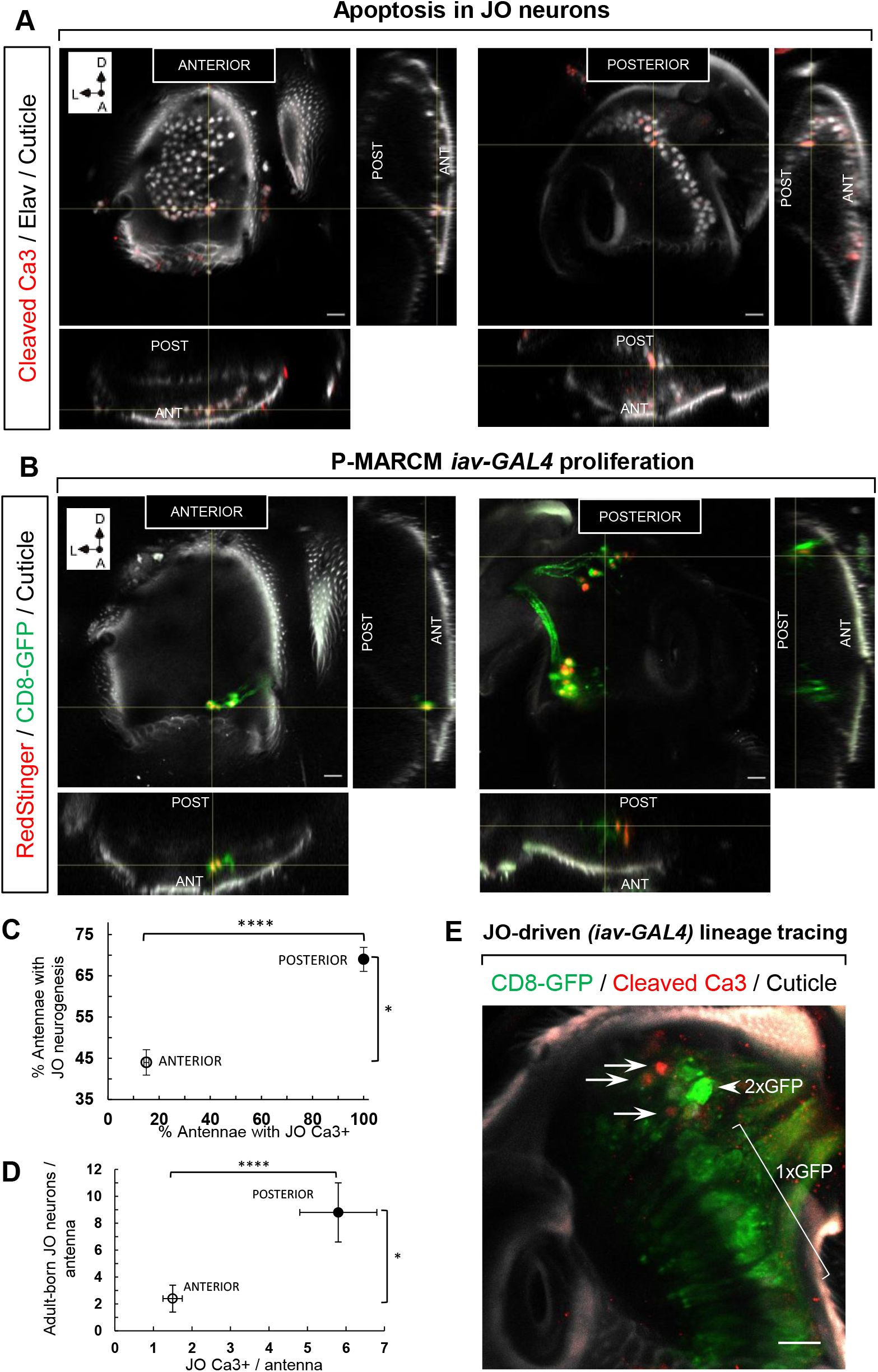
JO cell turnover occurs under physiological conditions. (A) Apoptotic JO neurons (Cleaved Ca3/Elav) are detected in anterior and posterior regions of the JO array. (B) P-MARCM *iav-GAL4* detects JO neurogenesis in posterior and anterior regions of the JO array. (C) The prevalence of adult JO neurogenesis as detected by P-MARCM (n=16 antennae) correlates with the frequency of apoptosis (n=33 antennae). Error bars represent s.e.m. \**p=0.012, ****p<0.0001*, cumulative probability on binomial distribution. (D) The number of adult-born JO neurons detected by P-MARCM (n=16 antennae) correlates with the extent of apoptotic JO neurons (n=33 antennae). Error bars represent s.e.m. *\*p=0.02, ****p<0.0001*, Mann-Whitney test. (E) JO neuron self-division (iav-GAL4, 2xGFP; arrowhead) occurs adjacent to apoptotic JO neurons (arrows). Non-dividing JO neurons contain a single ocpy of GFP (iav-GAL4, 1xGFP lineage tracing. Scale bars for all panels: 10μm.

### Enhancement of JO neuron regeneration through pharmacological administration

Turnover and regeneration of hair cells proceed at different extents in non-mammalian vertebrates (Bucks et al., 2017; Corwin and Cotanche, 1988; Ryals and Rubel, 1988; Williams and Holder, 2000). Since adult JO neurons turn-over under physiological conditions, we hypothesized JO neurons may also regenerate after different forms of injuries or under different environmental conditions.

Cisplatin is an agent used in cancer therapy with well-known ototoxic side effects: it induces hair cell death in vertebrates (Alam et al., 2000; Ou et al., 2007; Slattery and Warcho, 2010), followed by their limited regeneration (Mackenzie and Raible, 2012). In *Drosophila*, administration of cisplatin induces a decrease in the negative geotaxis behavior, a function mediated by JO neurons (Podratz et al., 2011). We therefore asked whether cisplatin-induced injury would increase JO neuron regeneration. We fed flies of the JO-driven GFP lineage tracing system with 50μg/ml cisplatin over 4 days (Podratz et al., 2011) and examined proliferation over the next 3 days, indicated by the presence of 2xGFP JO neurons (Fig 7A). Compared to 12% of control flies (n=26) exhibiting proliferation with 2.3 +/-1.1 s.e.m. 2xGFP new neurons/antenna on average (Fig. 7B, D), we identified an increase in proliferation in 28% of cisplatin-treated flies (n=18), with 4.6 +/-0.9 s.e.m. 2xGFP new neurons/antenna (Fig. 7C, D). Our results provide evidence for accelerated regenerative capacity of mechanosensory neurons in *Drosophila* following oral administration of this clinical anti-cancer compound.

**Figure 7.**
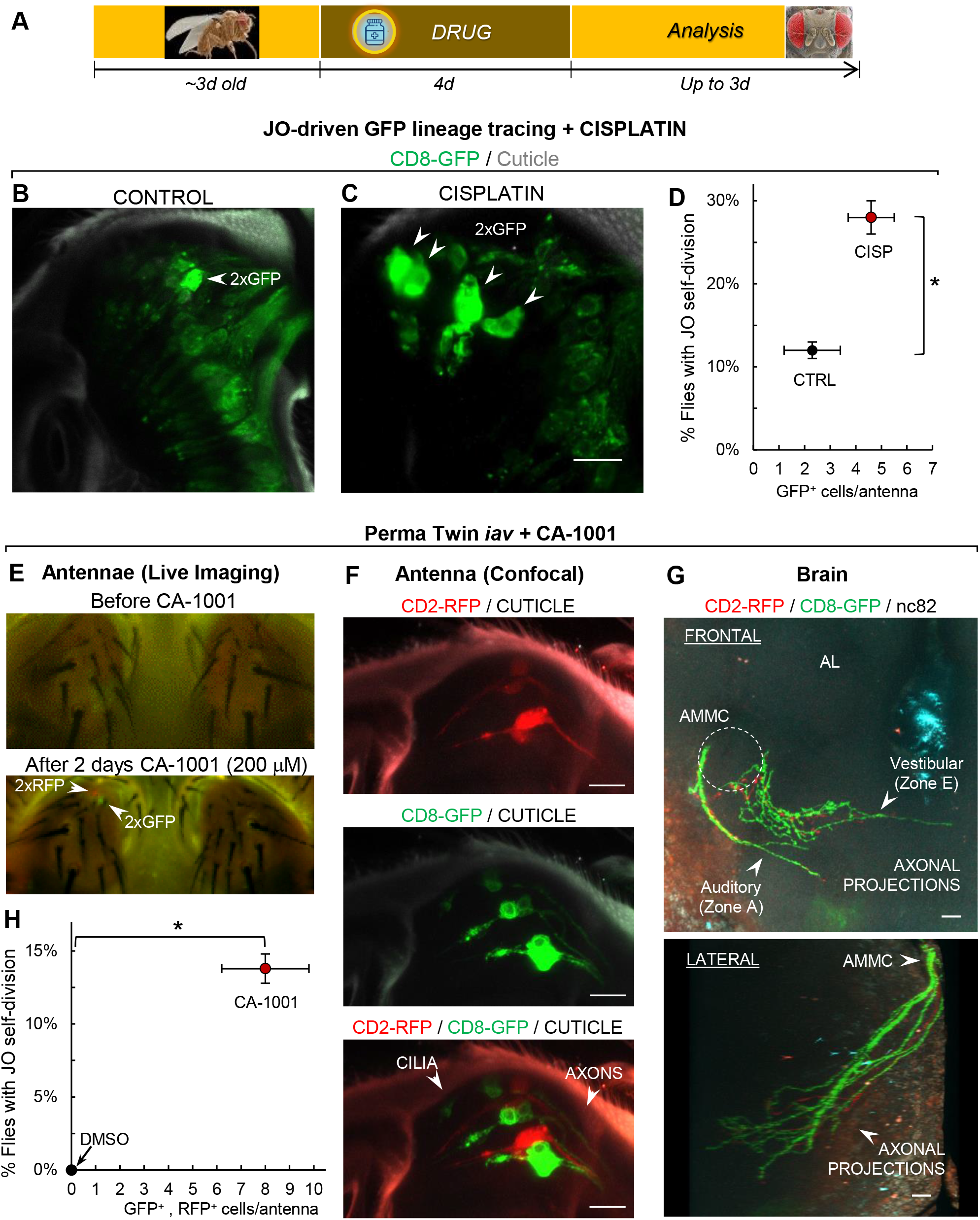
Enhancement of JO neuron regeneration through pharmacological administration. (A) Experimental timeline for pharmacological administration. Two-3 days-old flies were fed over 4 days with control DMSO or drug-supplemented food. JO neuron self-renewal was assessed up to 3 days later. (B-C) 2xGFP cells from JO-driven lineage-tracing reveals JO self-division in control (CTRL, B) and Cisplatin-treated (C) fly antennae. (D) Cisplatin administration increases the frequency and amount of JO neuron self-division (n=26 CTRL flies; n=18 cisplatin flies) *\*p=0.02*, cumulative probability of bionomial distribution. Bars represent s.e.m. (E) The calcium ionophore CA-1001 triggers self-renewal of JO neurons, as detected in live flies by Time-Lapse Imaging. New neurons are marked by the appearance of GFP+ and RFP+ hemiclusters (twins) using the PT-iav system. (F) New JO neurons develop cilia and send axons to the brain. Top panel: RFP hemicluster, Middle panel: GFP hemicluster, Bottom panel: Merged cluster. (G) CA-1001-induced JO neurons display Auditory (zone A) and Vestibular (zone E) circuit patterns. Axonal projections analyzed according to (Ishikawa and Kamikouchi, 2016). AL: Antennal Lobe; AMMC: Antennal Mechanosensory and Motor Center. Scale bars for all panels: 10μm. (H) CA-1001 administration increases the frequency and amount of JO neuron self-renewal (n=20 CTRL flies; n=29 CA-1001 flies). *\*p=0.02* by one sample t-Test. Bars represent s.e.m.

There is an increasing appreciation of the utility of *Drosophila* as an *in vivo* system for drug screening (Fernández-Hernández et al., 2016; Papanikolopoulou et al., 2019; Su, 2019). To expand our platform’s potential for identifying further compounds promoting the self-renewal of JO neurons, we also administered the drug CA-1001 to flies of the Perma Twin-iav system. CA-1001 is a calcium ionophore we identified in a previous small-molecule screening as a modulator of neurogenesis in the *Drosophila* central nervous system (Fernández-Hernández and Bonaguidi, unpublished data). By administering CA-1001 to Perma Twin-*iav* flies in a similar scheme as cisplatin (Fig. 7A), we observed JO neuron self-renewal using live time-lapse imaging. GFP+/RFP+ twins were detected in the antennae of intact adult flies as early as 2 days after the treatment was initiated (Fig. 7E). Confocal imaging revealed that new JO neurons develop cilia and project axons (Fig. 7F) to innervate the brain through the AMMC in both, the auditory (high frequency, Zone A) and vestibular (backward deflections, Zone E) circuits (Ishikawa and Kamikouchi, 2016) (Fig. 7G, Movie 1 and Fig. S6). Remarkably, we identified proliferation in 14% of the flies (n=20) with 8.0 +/-1.8 s.e.m. cells/antenna, compared to no proliferation detected in the DMSO-treated, control flies (Fig. 7H). Taken together, our results: i) demonstrate adult JO neurons have the capacity to respond to external stimuli (e.g., drugs) to adjust their self-renewal capacity; and ii) establish a new *in vivo* platform to screen for small molecules that accelerate the regeneration of mechanosensory cells.

## DISCUSSION

There is an urgent need to develop regenerative interventions for lost hair cells. However, the field has been hampered by the lack of *in vivo*, high-throughput platforms to easily assess the functional regeneration of adult sensory cells at the genetic, neural circuitry and behavioral levels. To address this challenge, we developed P-MARCM—a modified lineage tracing system in *Drosophila* for capturing cell type-specific proliferation in adult tissues over time without immunostaining. P-MARCM successfully detected the low-rate of neurogenesis and regeneration occurring in adult optic lobes, consistent with previous reports (Fernández-Hernández et al., 2013) (Fig. S1, S2). We leveraged the versatility of P-MARCM to identify adult genesis of JO neurons, the functional counterparts of vertebrate hair cells, by time-lapse imaging of intact flies and confocal microscopy. This platform enabled us to witness the addition of new cells in external organs *in vivo* over time, and to track cell proliferation at the single-fly level. Furthermore, the incorporation of an additional *UAS*-construct permits genetic manipulation of adult-born cells to assess their functional contributions. For example, future experiments could assess JO neurons’ contribution to auditory and vestibular function by applying existing behavioral protocols (Inagaki et al., 2010; Kamikouchi et al., 2009; Vaughan et al., 2014). Indeed, our results suggest new JO neurons have the potential to functionally modify mechanosensory circuitry, since they develop sensory cilia, express an essential mechano-transducer gene, and target appropriate brain regions for auditory and vestibular functions. This system will also enable research on mechanisms driving synapse formation between new and pre-existing neurons.

A central question in regenerative medicine is identifying a cell-of-origin that initiates tissue turnover. Analysis of P-MARCM images prompted us to hypothesize JO neurons could self-divide and remain in the tissue. Surprisingly, we identified low-level self-replication of JO neurons by live imaging using an independent JO-specific lineage tracing method. These results revealed the unexpected proliferative capacity of JO neurons, and confirmed P-MARCM results. We postulate that JO self-renewal can occur in *Drosophila* because: i) JO neurons are enclosed by 3 distinct supporting cells in the scolopidia, making it challenging to incorporate external JO neurons; ii) each scolopidia contains 2-3 JO neurons, a potential back-up mechanism to promptly replace lost JO without compromising other scolopidial cell types; and iii) self-division would facilitate proper cilia- and axon-targeting for optimal function in new JO. However, the differences in JO neurogenesis detected by JO-specific lineage tracing (20% of flies) compared to P-MARCM (45% of flies) suggest other cell types (undifferentiated- or supporting-cells activated in P-MARCM) might serve as progenitor cells. Alternatively, the observed differences could simply reflect recombination efficiencies between our methods. We further validated mitotic activity in JO neurons by a third lineage tracing system, PT-iav, which differentially labels only JO neurons produced through self-replication. Taken together, these 3 independent methods converge to demonstrate the generation of mechanosensory cells in the antennae of adult *Drosophila*, and to identify the self-renewal of JO neurons as an unexpected mechanism of sensory cell regeneration. Furthermore, these results rise the possibility that self-renewal of neurons might occur in other regions of the adult nervous system.

Recent efforts have established mammalian *in vitro* platforms for drug screening to promote hair cell renewal (Costa et al., 2015; Koehler et al., 2013; Koehler et al., 2017; Landegger et al., 2017). While useful, these platforms lack physiologic environmental and systemic cues, such as those controlling tissue interactions, as well as drug metabolism and availability. In our case, the low-levels of JO self-division and the sensitivity to identify newborn neurons by time-lapse imaging of individual flies provides an *in vivo* scalable platform to screen for small-molecules with translational relevance that enhance JO regeneration (Fernández-Hernández et al., 2016). Indeed, we provide a proof-of-principle of enhanced JO neuron regeneration upon the oral administration of cisplatin, a common ototoxic drug, and CA-1001, a calcium modulator. On one hand, cisplatin is known to kill hair cells in vertebrates, which regenerate afterwards up to a limited extent in the zebrafish lateral line (Mackenzie and Raible, 2012). Our results suggest a similar compensatory proliferation triggered in the JO neurons following cisplatin administration. On the other hand, CA-1001 is a calcium ionophore which facilitates the transport of Ca2+ across the plasma membrane. Here, Ca^2+^ might play a direct role in the self-division of JO neurons, since calcium signaling is a known regulator of cell proliferation, the expression of genes involved in cell growth, and the early steps of neurogenesis (Leclerc et al., 2012; Pinto et al., 2015; Resende et al., 2013). Although elucidation of the molecular mechanisms driving regeneration of JO neurons is pending, for these as well as for other selected drug hits, the transcriptomic and epigenetic changes in JO neurons can be assessed as they occur *in vivo*, by available genetically-encoded tools (Marshall and Brand, 2017; Marshall et al., 2016; Southall et al., 2013). Further, the functional contribution of regenerated JO can be readily assessed by established behavioral protocols (Kamikouchi et al., 2009; Sun et al., 2009; Vaughan et al., 2014). In summary, the new *Drosophila* platform, presented here, represents a promising approach to identify modifiers of neuronal regeneration, their mechanisms of action and their functional consequences at both the circuitry and behavioral levels.

## MATERIALS AND METHODS

### Fly lines and experimental conditions

#### For P-MARCM-iav and MARCM-iav experiments

Female virgins of genotype *hs-Flp,tub-GAL80,neoFRT19A; hs-FlpD5,20UAS-6GFPmyr, UAS-RedStinger / CyO; iav-GAL4* were crossed to males of genotype *tub FRT STOP FRT lexA,neoFRT19A; +; 8lexAOp-Flp/TM6B*. The cross was set and kept at 17°C during development to minimize spontaneous *hs-Flp* activation. For P-MARCM, we picked 2-5 days old female virgins with final genotype: *hs-Flp,tub-GAL80,neoFRT19A / tub FRTSTOP FRTlexA,neoFRT19A; hs-FlpD5,20UAS-6GFPmyr,UAS-RedStinger / +; iav-GAL4 / 8lexAOp-Flp*. For MARCM, females of same age and genotype, but carrying a *TM6B* balancer chromosome instead of *8lexAOp-Flp* were picked. *hs-FlpD5* (Nern et al., 2011) was inserted to maximize FLP induction and recombination. Every single fly of these genotypes was then pre-screened under an epi-fluorescent scope to ensure only those with minimum-to no-background labelling were used for proliferation analysis. Selected flies were then pooled, and control ones randomly picked for dissection corresponding to “0w” time-point. Remaining flies were allocated for activation by heat shock and dissection at later time points. In order to maximize the number of cells with the system activated, flies were heat-shocked at 38°C (Ohlstein and Spradling, 2006) for 30 minutes twice on the same day, ~2 hours apart.

#### For P-MARCM-nsyb experiments (Fig. S1)

Here, flies of final genotype *hs-Flp,tub-GAL80,neoFRT19A / tub FRT STOP FRT lexA,neoFRT19A; 20UAS-6GFPmyr,UAS-RedStinger / +; nsyb-GAL4 / 8lexAOp-Flp* were used. Since preliminary assessment of background labeling in the brain is not possible without dissection, only 1 copy of regular *hs-Flp* was included to minimize leaky expression. Flies were blindly assigned to the control group for dissection and to the experimental group for heat shock at 38°C for 45 minutes twice on the same day, ~2 hours apart, and dissected 3w later.

#### For JO-driven (iav-GAL4) lineage tracing system

Female virgins of genotype *w; UAS-CD8-GFP,UAS-CD2mir, FRT40A; 20UAS-FlpD5* were crossed to males of genotype *w; tubQS,FRT40A; iav-GAL4*. We selected males and females, 2-5 days old, with the final genotype *w; UAS-CD8-GFP,UAS-CD2mir,FRT40A /tub-QS,FRT40A; 20UAS-FlpD5 / iav-GAL4*. These flies were then screened individually by fluorescent microscopy to remove those with background labeling. Selected flies were tracked weekly over 4 weeks, and those with JO neurogenesis were live-imaged by fluorescent microscopy (see below) and harvested for confocal imaging, as described below.

#### For PT-iav lineage tracing system

Female virgins of genotype *w; UAS-CD8-GFP,UAS-CD2mir,FRT40A; 20UAS-FlpD5* were crossed to males of genotype *w; UAS-CD2-RFP,UAS-GFPmir,FRT40A; iav-GAL4*. Males and females, 2-5 days old, with the final genotype *w; UAS-CD8-GFP,UAS-CD2mir,FRT40A / UAS-CD2-RFP,UAS-GFPmir,FRT40A; 20UAS-FlpD5 / iav-GAL4*. These flies were screened individually by fluorescent microscopy to remove those with background labeling. Selected flies were either tracked weekly between 1 and 4 weeks in physiologic conditions or up to 3 days after drug administrations for JO neurons regeneration.

### Drug treatment

Cisplatin (Sigma, 479306-1) was freshly prepared dissolved at 50μg/ml (final concentration) in ddH2O. As a control, ddH2O was used. CA-1001 (Cayman, 17407) was prepared as a 10 mM stock solution in pure DMSO at and stored at −20C, for further dilution at 100μM or 200μM for experimental use. As control, DMSO 1% or 2% were used. About 200 mg of instant fly food in powder form was reconstituted with 1ml of the corresponding drug or control solution in an empty plastic vial, where <10 flies were placed for treatment over 2 days, when they were transferred to another vial with fresh drug-containing food for an additional 2 days. After a total 4-days treatment, flies were transferred to vials with regular food for tracking JO neurogenesis up to 3 days later. Those with JO neurogenesis were live-imaged by fluorescent microscopy (see below) and harvested for confocal imaging as described below.

#### Dissection, immunostaining, and confocal imaging

Antennae were dissected, attached to their corresponding brains in chilled Schneider’s medium, and then fixed in 3.7% formaldehyde solution for 20 min. They were then washed with PBT 1% solution for 10-20 min, followed by a final wash in 1XPBS before incubation with primary antibodies overnight at 4C (2 nights for nc-82 antibody), followed by incubation with secondary antibody for 4 hours at room temperature or overnight at 4C. For staining of neurons and caspase, the third segment of antennae was removed before fixation, to facilitate antibody penetration to JO neurons. Primary antibodies were: mouse anti-nc82 (1:10, DSHB deposited by Buchner, E. (Wagh et al., 2006), rat anti-elav (1:100, DSHB, deposited by Rubin, Gerald M. (O’Neill et al., 1994), rabbit anti-cleavedCaspase3 (1:200, Cell Signaling); secondary antibodies (Jackson laboratories) were: anti-mouse Cy5 antibody (1:100), anti-rat Cy5 (1:100), anti-rabbit Cy3 (1:200), and anti-rabbit Cy2 (1:200). No antibodies were used for GFP, RFP, or RedStinger fluorescent proteins. Antennae and brains were mounted in Vectashiled media with DAPI (Vector laboratories). For mounting, we used double-side sticker spacers (EMS, 70327-9S) to preserve morphology as much as possible. We used 1 spacer for antennae and 2 for matching brains. Images were acquired on a Zeiss LSM 700 confocal microscope at either 1.2 μm spacing with 20X objective or 0.6 μm spacing with 40X objective. Images shown represent maximum intensity projections of relevant planes or 3D projections.

#### Time lapse imaging of live flies

Flies were anesthetized on a CO2 pad for imaging under a Zeiss V16 epi-fluorescent scope with a 5.6Mp monochrome camera. These acquisition settings were selected to image P-MARCM flies in order to preserve viability after imaging over time: 50% power lamp, PlanApo 1X objective, 85% aperture, 160X total magnification, 5×5 camera binning, 90ms exposure time for GFP and RFP channels, and, on average, a ~65 um Z-stack with 4um increments. For JO-lineage tracing 4×4 camera binning was selected. Maximum intensity projections were generated for display.

#### Gaussian mixture model method

Labeled neurons were counted in the JO of each fly, from all time points. Assuming JO from any time point fell into either “Responder” or “Non-Responder” categories, we fit a Gaussian mixture model with 2 mixture components using the mclust package (Scrucca et al., 2016) with default parameters. The model found that any fly with 9 or fewer neurons was in the Non-Responder category (*i.e.*, comparable to system background levels), accounted for 63% of all observations, while any fly with 10 or more neurons was a Responder, (*i.e.*, above background levels) accounted for the remaining 37% of all observations.

### Statistical analysis

Plots and statistical analysis were done using GraphPad Prism and Microsoft Excel with data from at least 3 replicate groups (2 groups for cisplatin tests), based on Student’s t-test or the cumulative probability of binomial distributions. s.e.m. on experiments with binary outcomes was calculated as 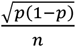 where *p* is the frequency of JO neurogenesis in the analyzed group and *n* the number of flies considered. In the quantification of JO neurons labeled with MARCM and P-MARCM systems (Fig. S4), any value above 2.5 s.d. from the mean was considered an outlier. This yielded for MARCM: 1/33 outlier at 0w (28 JO neurons labeled); and for P-MARCM: 1/17 outlier in 0w group (45 JO neurons labeled) and 1/19 outlier in the 1w group (42 JO neurons labeled), which were excluded from analysis.

## ACKNOWLEDGEMENTS

We thank: members of the Bonaguidi lab for support, especially Maxwell Bay for assistance with the Gaussian model and statistical analysis, and Eric Hu for technical assistance with maintenance of fly lines; Dr. Neil Segil (USC) for directing our attention to the auditory system of the fly and for insightful discussions on this project; Dr. Matthias Landgraf (U. Cambridge) for kindly providing *tub>STOP>lexA* flies; Cristy Lytal for editing the manuscript; Bloomington Drosophila Stock Center (National Institutes of Health P40OD018537) for other lines used in this study; and Developmental Studies Hybridoma Bank, created by the National Institute of Child Health and Human Development of the National Institutes of Health for nc-82 and elav antibodies.

## AUTHOR’S CONTRIBUTIONS

**Ismael Fernández-Hernández**

Conceptualization; methodology; investigation; formal analysis; writing (original draft); funding acquisition

**Evan B. Marsh**

Investigation

**Michael A. Bonaguidi**

Conceptualization; resources; writing (review and editing); supervision; funding acquisition

## COMPETING INTERESTS

The authors declare no competing interests.

## FUNDING

Authors acknowledge support from USC-CONACYT (Consejo Nacional de Ciencia y Tecnología) Postdoctoral Scholars Program Fellowship and USC Provost’s Postdoctoral Scholar Research Grant to I.F.-H.; USC Provost Undergraduate Research Fellowship to E.B.M.; and National Institutes of Health (R00NS089013, R56AG064077), Whittier Foundation, Baxter Foundation and Eli and Edythe Broad Foundation grants to M.A.B.

---

**Fig. S1.**
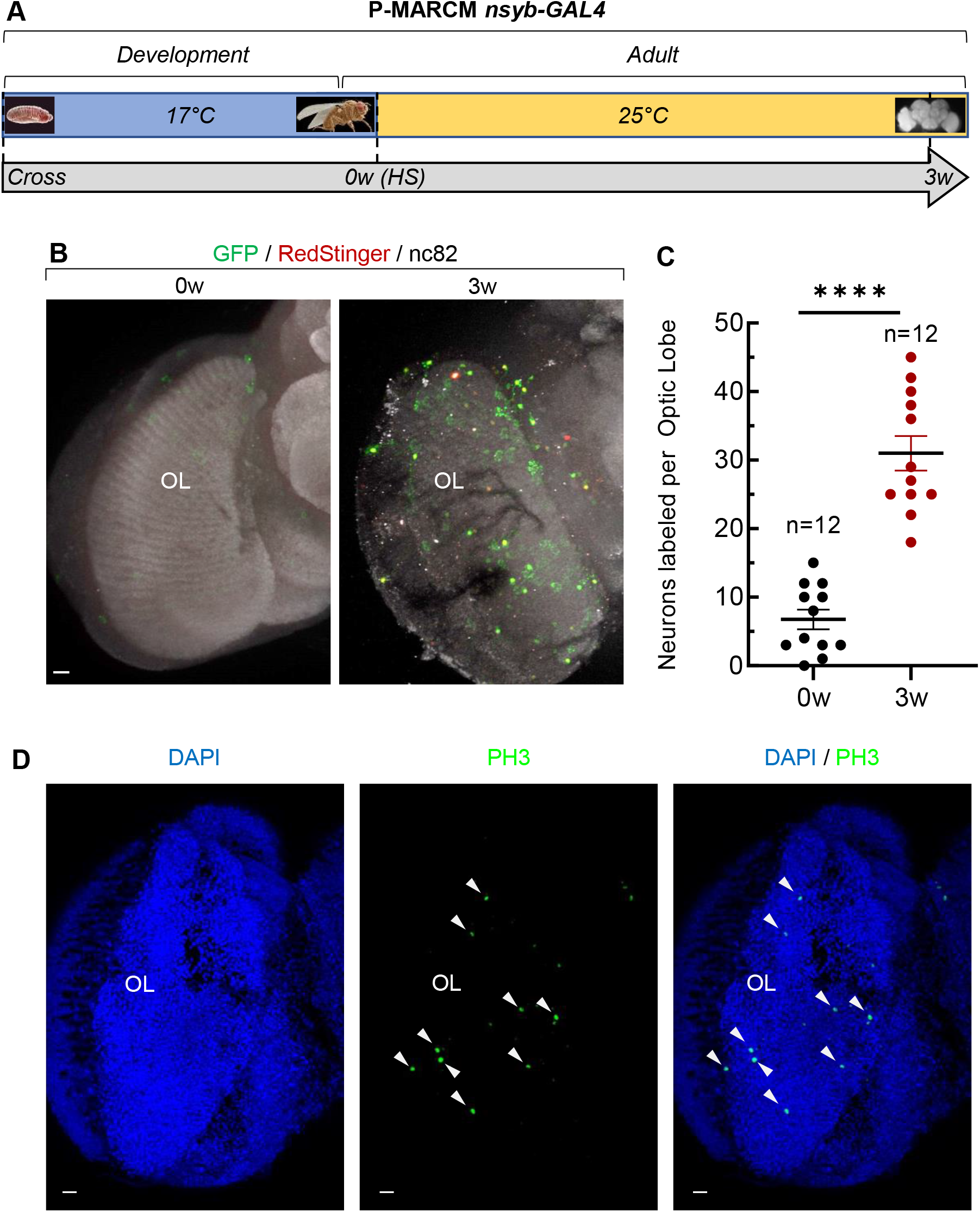
P-MARCM captures adult neurogenesis in the *Drosophila* optic lobes (OL) (A) Experimental strategy to reveal adult neurogenesis with P-MARCM *nsyb-GAL4*. Flies 2-5 days–old were Heat-Shocked (HS) to activate the P-MARCM system and brains were dissected 3 weeks (3w) after. (B) Adult-born neurons in the optic lobes (OL) are labeled by P-MARCM with *nsyb-GAL4* line 3 weeks after HS. (C) Amount of adult-born neurons in OL at 3 weeks (n=12 OL) is significantly higher than background levels (n=12 OL) (*p=0.0000002*, Student’s t-test). Error bars represent s.e.m. (D) Cell proliferation is also detected by anti-PH3 antibody in the OL (9.2+/-1,0 s.e.m. PH3+ cells/OL; n=26 OL). Scale bars for all panels: 10mm.

**Fig. S2.**
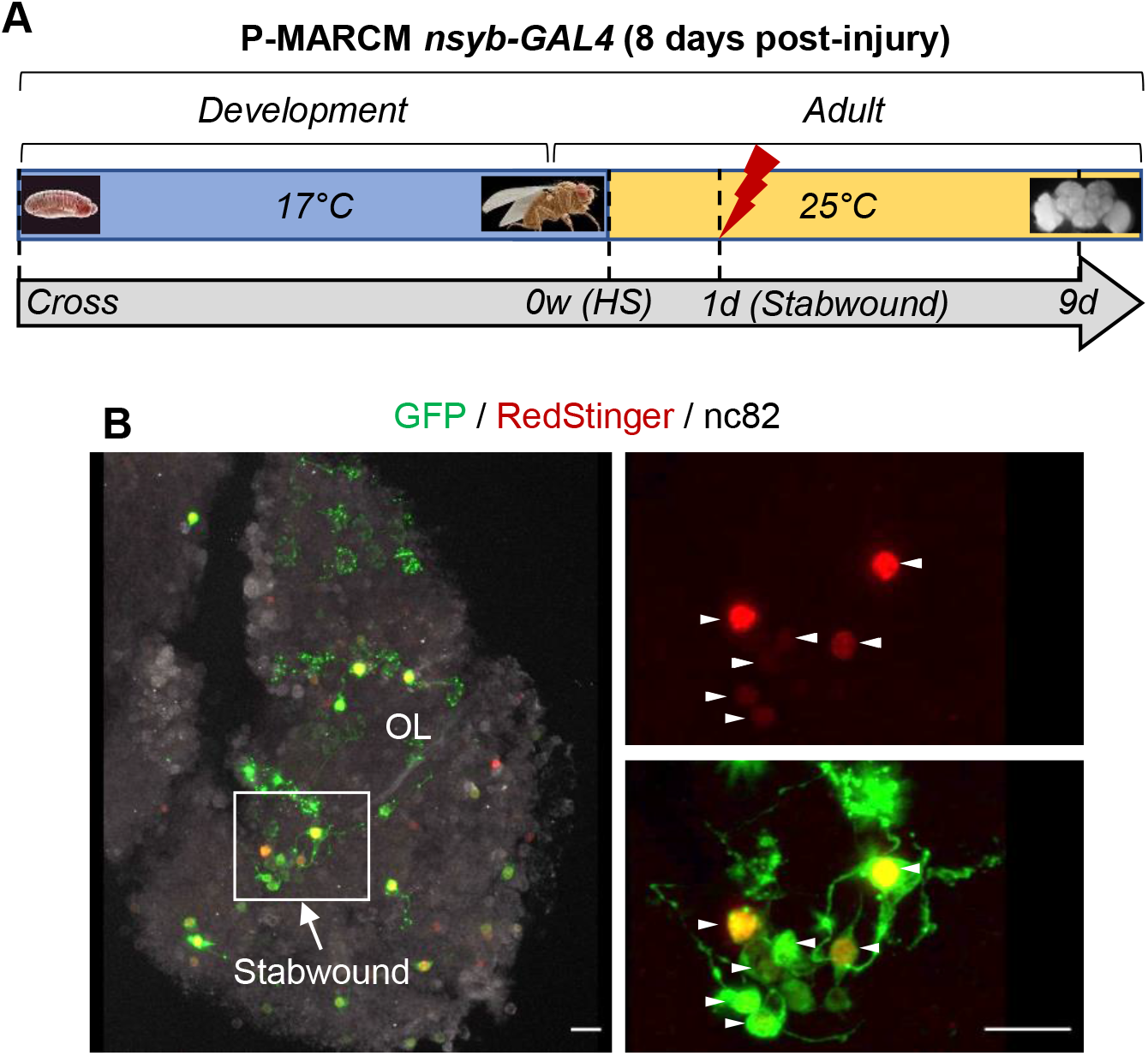
P-MARCM captures injury-induced neuronal regeneration in the *Drosophila* optic lobe (OL) (A) Experimental strategy to capture injury-induced neuronal regeneration in OL with P-MARCM *nsyb-GAL4*. Two to 5 days–old flies were Heat-Shocked (HS) to activate the P-MARCM system. Stab wound was applied to the left OL by a fine needle 1 day after HS. Flies were dissected and imaged 9 days later. (B) Regenerated neurons in the OL (arrowheads) are labeled by P-MARCM with *nsyb-GAL4* line 9 days after stab wound. Scale bar: 10 μm.

**Fig. S3.**
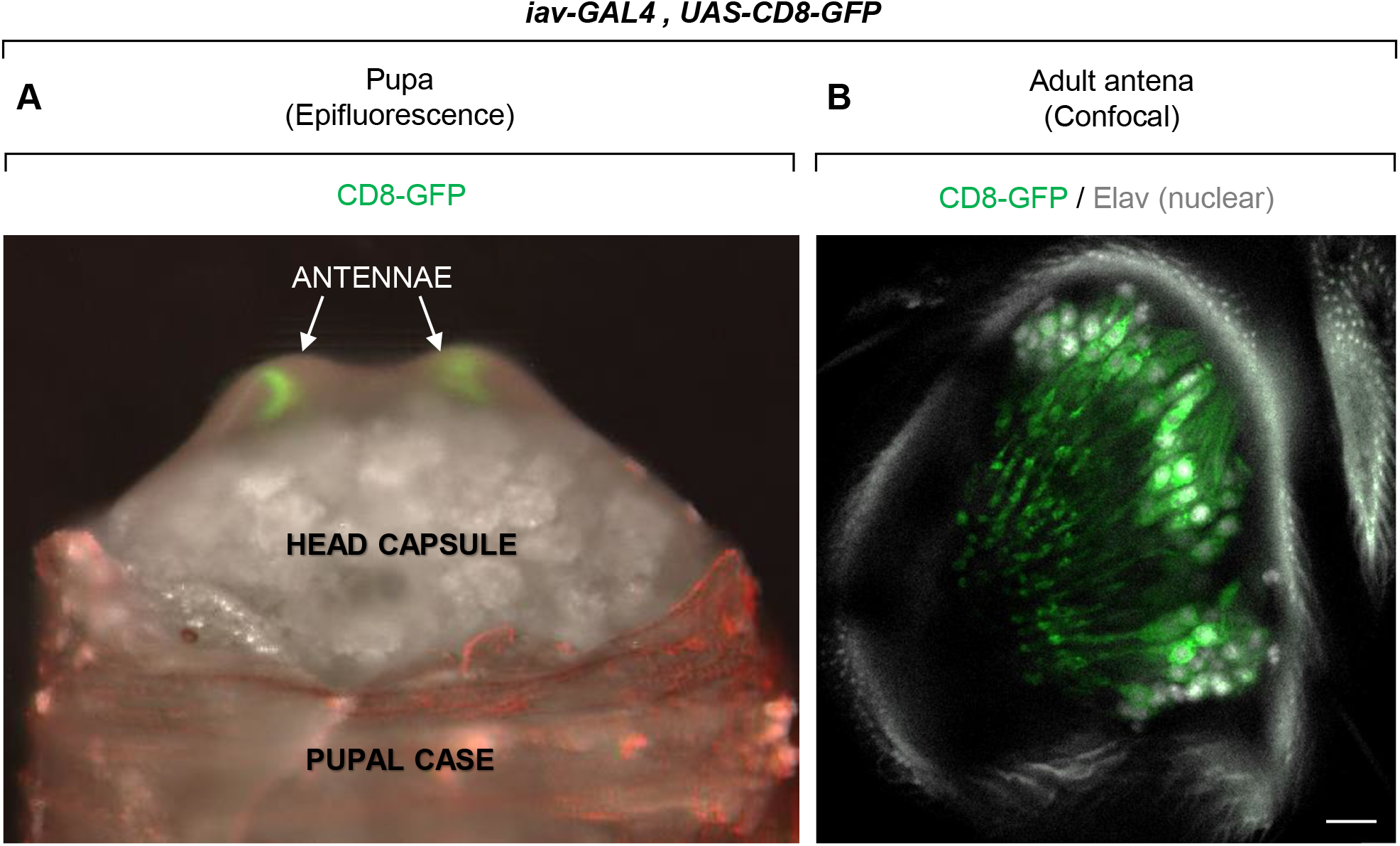
Expression pattern of *iav-GAL4* line in the antennae. (A) Expression of *iav-GAL4* in the antennae begins in pupal stage. (B) *iav-GAL4* expression is restricted to JO neurons in adult antenna. Scale bar: 10μm.

**Fig. S4.**
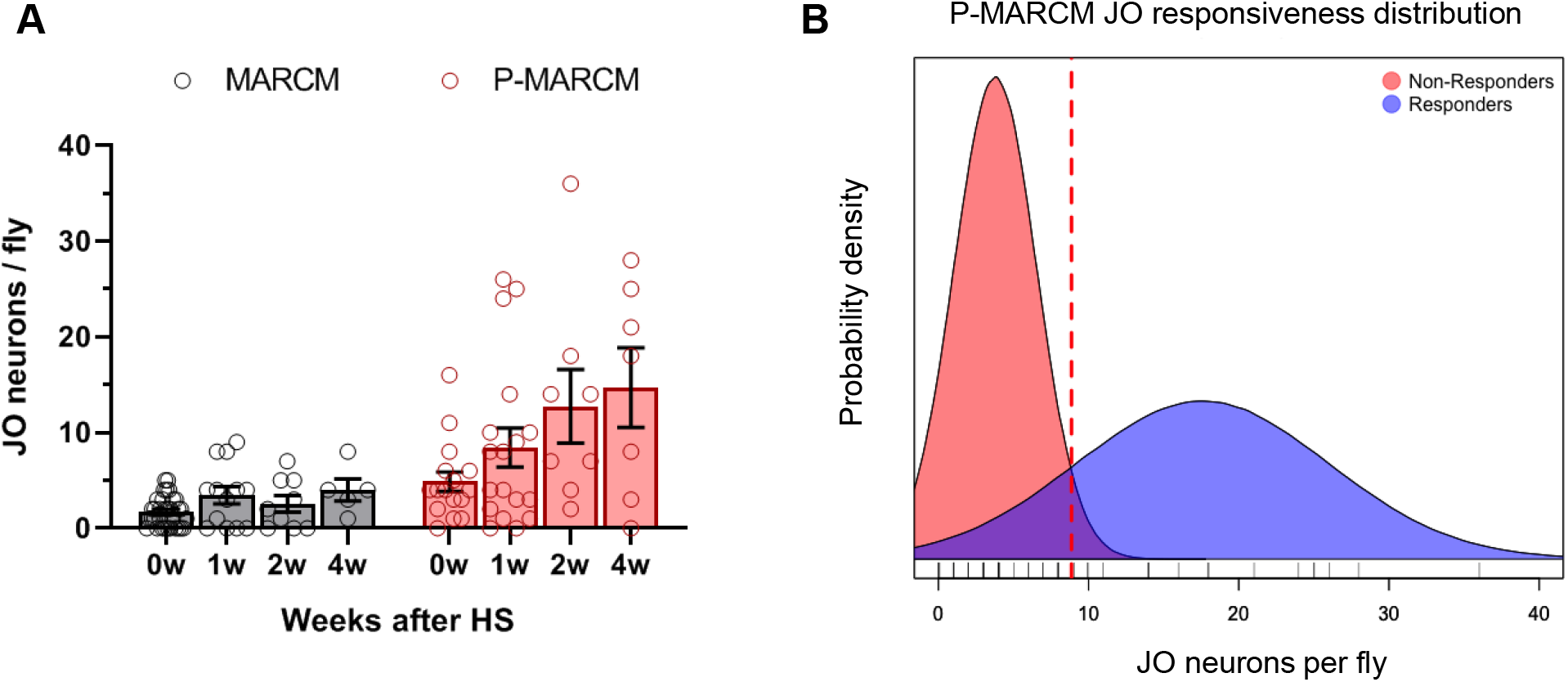
JO neurogenesis detection and distribution. (A) Quantification of newborn JO neurons in MARCM and P-MARCM lineage tracing approaches. JO neurogenesis increases over time upon P-MARCM labeling. Error bars represent s.e.m. (B) Gaussian mixture model classifies flies into “Responders” and “Non-responders” indicating the presence of adult JO neurogenesis beyond background levels.

**Fig. S5.**
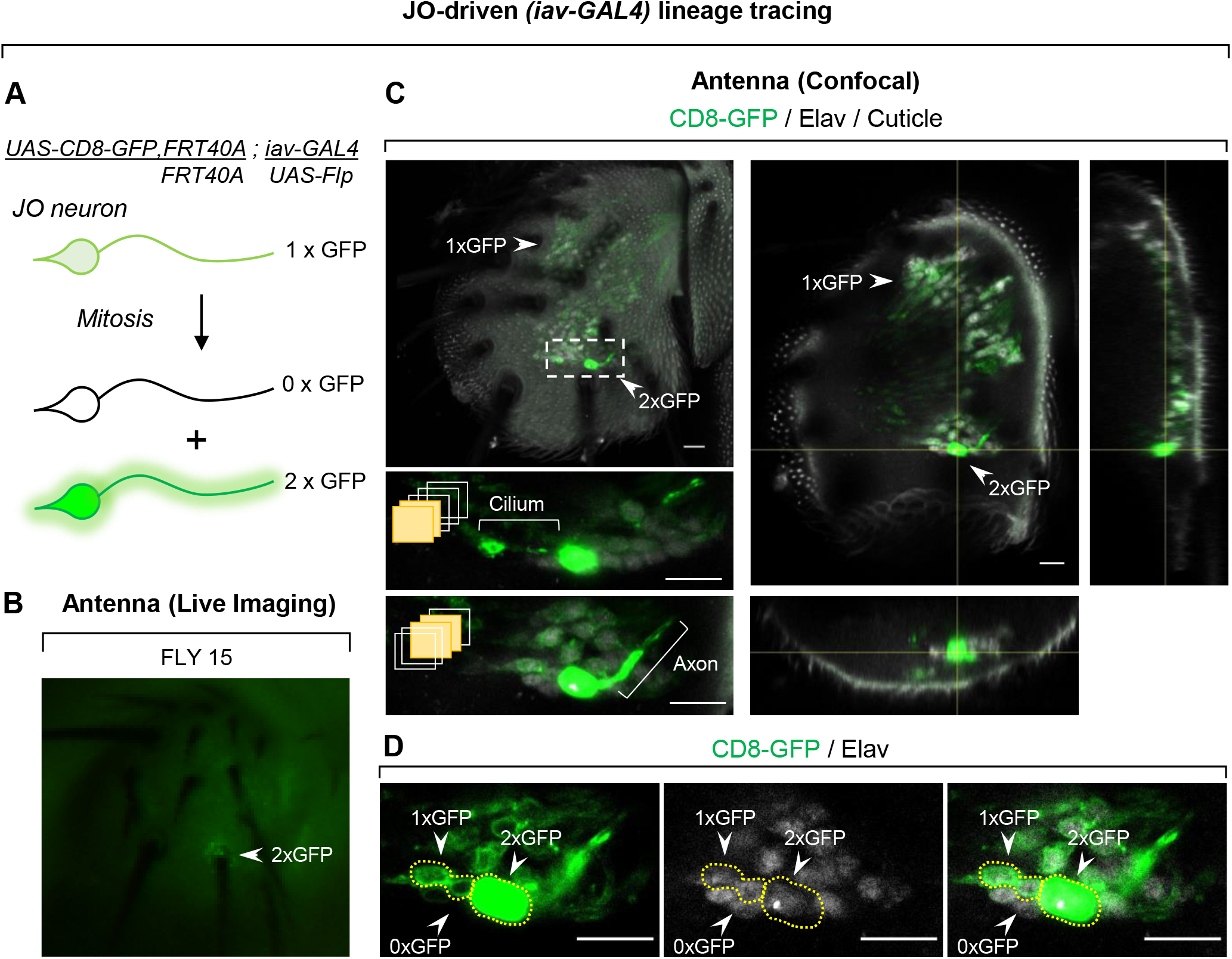
Self-dividing JO neurons are captured at single-cell resolution *in vivo*. (A) *.iav-GAL4*-driven lineage tracing system to assess JO self-division. Daughter cells express 2xGFP and 0xGFP, while non-dividing JO neurons contain 1xGFP. (B) A single JO neuron expressing 2xGFP detected by live Time Lapse Imaging. (C) Confocal microscopy confirms proliferation of a single neuron on the anterior part of the antenna. Captions show maximum intensity projections of discrete planes to visualize the JO neuron cilium and axon. (D) JO neurons self-division detected by twin-spots of 2xGFP/Elav+ neurons and 0xGFP/Elav+ neurons among non-dividing 1xGFP/Elav+ JO neurons. Scale bars: 10μm

**Fig. S6.**
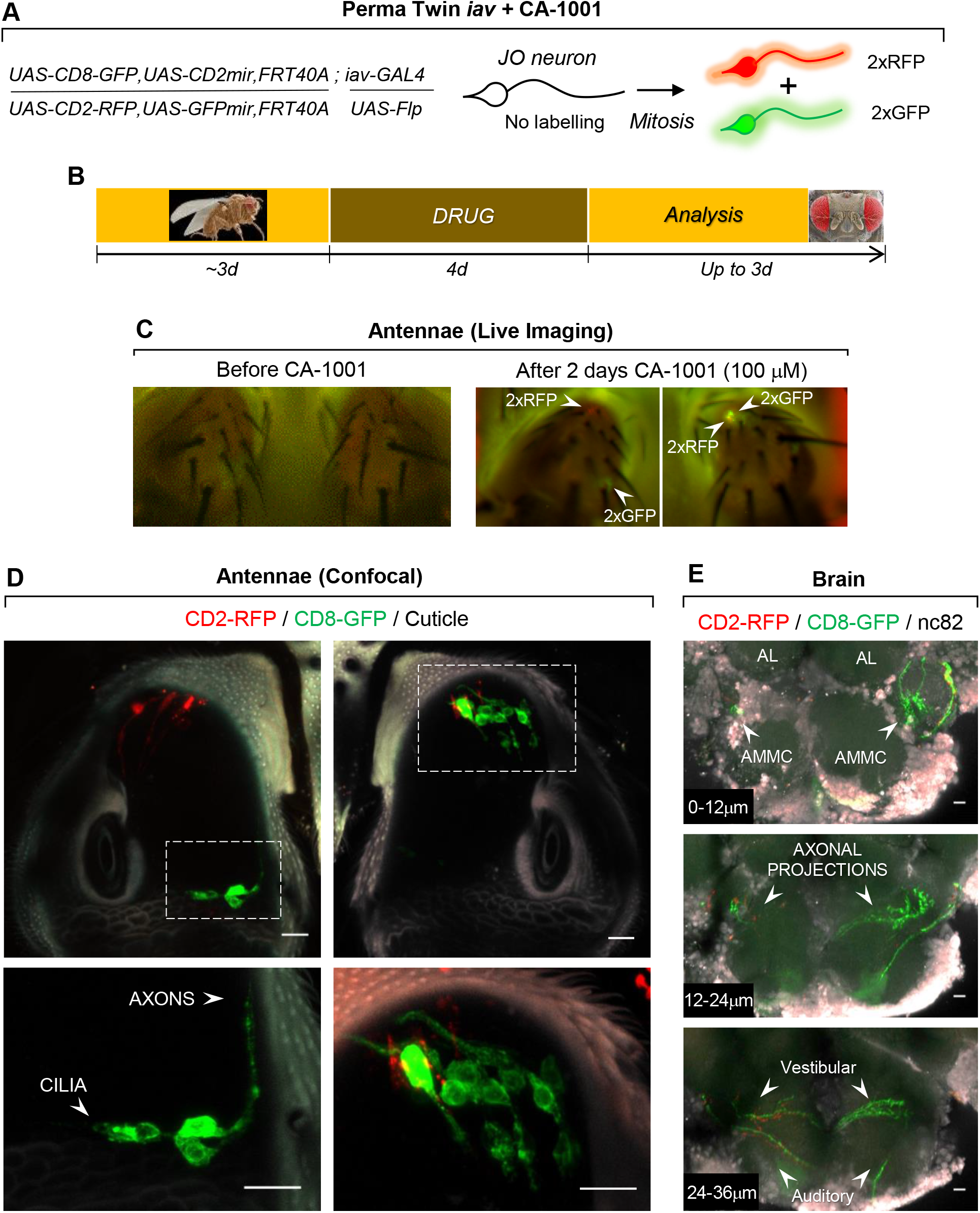
The calcium ionophore CA-1001 increases JO neuron self-renewal. (A) Perma Twin-iav lineage tracing system to assess JO neuron self-replication. (B) Experimental strategy to capture JO neuron self-renewal. Three-day old PT-iav flies receive oral drug administration for 4 days and are analyzed up to 3 days later. (C) Self-replicated JO neurons are detected by live Time Lapse Imaging as soon as 2 days after administration of CA-1001 at 100μM. (D) Self-replicated JO neurons develop cilia and extend axons to the brain. AMMC: Antennal Mechanosensory and Motor Center; AL: Antennal Lobe (E) New JO neurons target the brain in the Auditory and Vestibular circuit pattern as early as 2 days following drug administration. Scale bars for all panels: 10μm.

## Notes

### Competing Interest Statement

The authors have declared no competing interest.

### Summary of Updates

Self-replication of JO neurons shown by an additional system. Regeneration of JO neurons is enhanced upon oral administration of drugs.

